# Plasticity in the pectoral fin skeleton is induced by altered foraging regime in a South American cichlid

**DOI:** 10.1101/2024.09.24.614498

**Authors:** Michelle Gilbert, Alexandria J. Kwiatkowski, Ciara M. Woodburn, Sofia N. Piggott, Yara Haridy, Brett R. Aiello, R. Craig Albertson, Thomas A. Stewart

**Affiliations:** Department of Biology, The Pennsylvania State University, University Park, PA 16802, USA; Biology Department, University of Massachusetts, Amherst, MA 01003, USA; Ecology, Evolution, and Organismal Biology Graduate Program, Brown University, Providence, RI, 02912, USA; Department of Organismal Biology and Anatomy, The University of Chicago, IL 60637, USA; Department of Biological Science, Seton Hill University, Greensburg, PA, 15601, USA

**Keywords:** Adaptation, Plasticity, Constraint, Phenotype, Skeleton, Swimming

## Abstract

The fins of fishes are remarkably diverse, and this variation is tied to the ecology and locomotor mode of a species. While numerous genetic factors are known to pattern fins in development, it is unclear how developmental plasticity shapes the fin skeleton. Here, we analyze the cichlid *Satanoperca daemon*, raised under three distinct feeding regimes, and show that plasticity is pervasive across the pectoral fin skeleton with foraging mode impacting patterning of both the endoskeleton and dermal skeleton. Radials and fin rays were µCT scanned and analyzed using a combination of linear measures and geometric morphometrics. Anteroposterior patterning of both radials and fin rays are affected by feeding regime. Notably, *S. daemon* pectoral fin rays show distinct patterns of fin ray branching between treatments, suggesting altered fin stiffness. We argue that the observed changes in the fin likely reflect developmental plasticity resultant from altered swimming behaviors when fishes are challenged to forage in different ways. These data show how non-genetic mechanisms can shape both the endoskeleton and dermal skeleton of fins, and that foraging mode can induce plastic changes in skeletal elements that do not directly interface with food items.

## INTRODUCTION

Studies of developmental plasticity can aid in the identification of traits from which novel variation is likely to arise as well as those that are resistant to developmental perturbations, thereby informing the likely direction of phenotypic evolution [1–8][9–11]. The ray-finned fishes (Actinopterygii) make up more than half of vertebrate species diversity [12], and the evolution of their disparate fin morphologies was likely key to the group’s evolutionary success. How developmental plasticity might contribute to fin disparity in actinopterygians is an outstanding question, as it is unclear which fin traits are plastic and what perturbations can impact fin development.

Fish use their fins for myriad functions, including protection [13], display [14], sensing [15–20] and, critically, locomotion [18,21–24]. In actinopterygians, fins comprise a proximal endoskeleton (radials) and a distal dermal skeleton (fin rays) [25]. Fin rays are bilaminar, composed of two hemitrichia, can be segmented and branching, and have muscles that attach at the base to control their curvature and stiffness [26]. The morphology of a fin is predicted to correspond to its functions [18,20–23,27,28] and, thus, explaining the developmental mechanisms that pattern the endoskeleton and dermal skeleton are key to explaining the radiation of fishes. While there have been great strides in understanding the genetic basis of fin patterning, development, and diversity [29–36], significantly less is known regarding the role of plasticity in fin evolution.

Several musculoskeletal features of the fin have been shown to be plastic, but which properties of fins can respond and under what conditions has not been well characterized. In *Polypterus senegalus*, plasticity of the pectoral fin skeleton was induced by rearing juveniles under semi-terrestrial conditions. Following treatment, Du and Standen [37] observed altered shape and size of proximal radials and number of distal radials in the pectoral fins. In cichlids, Navon and colleagues reported that forcing juveniles to forage in different ways resulted in alterations to the pectoral fins, including altered fin ray number and shape of fin musculature, suggesting that pectoral fin morphology could reflect feeding behaviors [38]. In the catfish *Phractocephalus hemioliopterus*, induction of the fin rays in the adipose fin appears plastic; captive individuals lack adipose fin rays, which are uniformly present in large wild-caught individuals [39]. Collectively, these studies implicate developmental plasticity as a mechanism capable of affecting fin architecture, altering the number and shape of fin skeletal elements and potentially contributing to major ecological transitions, including the water-to-land transition of early tetrapods [37,40,41]. However, it is unknown whether fin ray morphology (*e.g.*, patterns of branching, cross-sectional morphology, or segment length) can be impacted, as well as whether plasticity can affect patterning of the pectoral fin skeleton along its various axes.

Cichlids have emerged as robust models for testing hypotheses of plasticity in vertebrate evolution due to their behavioral flexibility and amenability to laboratory-based experimentation [42,43]. Previously, the South American cichlid *Satanoperca daemon* was leveraged for understanding craniofacial plasticity [44] due to its unique winnowing behavior. This foraging behavior involves the oral intake of sediment, the mechanical sifting of food (plant detritus and invertebrates) from the substrate, and ends with the expulsion of the sediment either past the gill plate or orally. Since *S. daemon* use their pectoral fins in maneuvering and in positioning during feeding, we tested whether endo- and dermal-skeletal components of the pectoral fin were affected when fish were induced to feed from different substrates. Here we report that altered foraging regimes can induce plastic responses in pectoral fin, affecting patterning and canalization of the fin skeleton. These results demonstrate that the environment and behavior of fishes can drive alterations across the fin which has implications for evolution and diversification of fins broadly.

## METHODS

### Experimental design

Wild-caught juvenile *Satanoperca daemon* were raised under three feeding regimes. All individuals were fed bloodworms but induced to forage in different ways by altering the tank environment (Figure 1*a*, Supplementary Movies 1-3). They were raised: (1) in tanks without substrate, which prevented winnowing, where fish foraged from food in the water column (n = 11); (2) in tanks requiring foraging by winnowing on fine sand (n = 14); (3) in tanks that required foraging by winnowing on small aquarium rocks (n = 9). We refer to these treatments as pelagic, sand-winnowing, and rock-winnowing treatments, respectively. One specimen in the rock-winnowing treatment developed a pigmentation pattern as a juvenile that suggests an identity of *Satanoperca jurupari*. In both past study of the craniofacial skeleton [44] and in this analysis, the individual’s morphology falls within the distribution of *S. daemon* for the rock-winnowing treatment, and so it was kept in the analysis.

**Figure 1.**
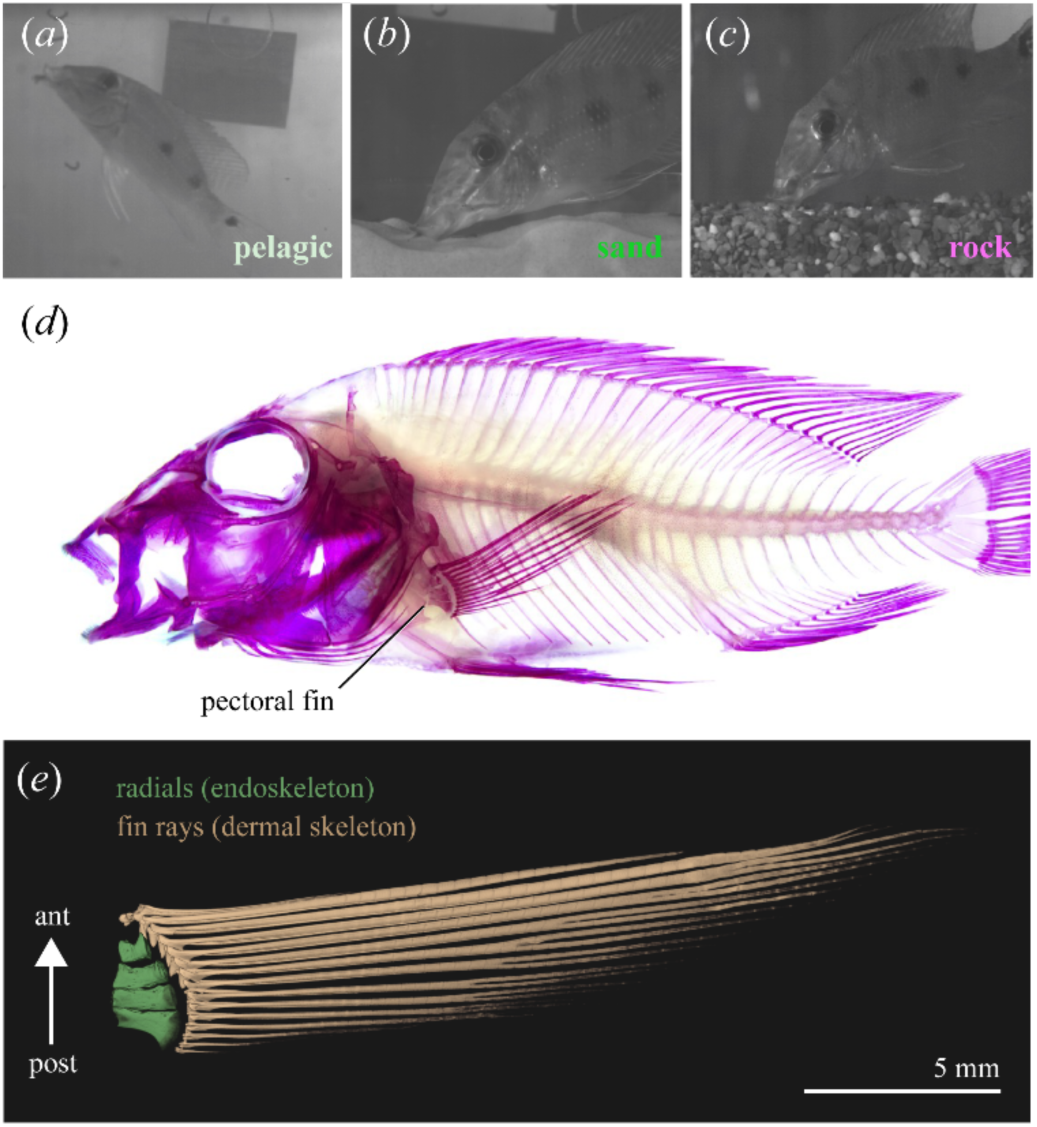
Juvenile *S. daemon* raised under three experimental treatments. (*a*) *S. daemon* feeding in the pelagic treatment, in which individuals were induced to feed in the open water. (*b*) *S. daemon* feeding in the sand-winnowing treatment. (*c*) *S. daemon* feeding in the rock-winnowing treatment. Following five months of treatment, pectoral fin skeletons were analyzed to test for plasticity. Footage of *S. daemon* feeding experiments is provided in Supplementary Videos 1-3. (*d*) Cleared and stained *S. daemon*. (*e*) Volumetric rending of µCT scan of a left pectoral fin showing the endoskeleton and dermal skeleton. (Abbreviations: ant, anterior; post, posterior)

Following five months of experimental treatment, specimens were euthanized (MS-222 and ice shock combination) and then fixed in 4% paraformaldehyde. All experimentation was performed in accordance with animal care protocols of the University of Massachusetts, Amherst. Additional details on specimen acquisition and the experimental design are provided in Gilbert et al 2022 [44].

To analyze the pectoral fin skeleton, fixed fish were stained using a 75% EtOH alizarin solution to stain bone while preserving muscle (Figure 1*b*). The left pectoral fin was dissected, and skin and musculature were removed to allow better visualization of radials and fin rays. Images that included a superimposed scale bar of each fin were taken using a Zeiss Discovery V20 and a Axiocam 506 Mono camera. Dissected pectoral fins were then micro-computed tomography scanned (µCT), described below (Figure 1*e*). Traits that could be assessed from external morphology were assessed on both left and right pectoral fins, and those traits that required dissection were analyzed for the left pectoral fin only.

### Counts and linear measures

For each fin, we counted the number of fin rays and collected linear measurements from photographs in ImageJ (v1.54h) [45]. Specifically, to characterize the endoskeleton, we measured length and width of radials from the dissected left side of the specimen, and to characterize the dermal skeleton, we measured the length of the first segment (*i.e.*, the long proximal segment) of fin rays 3-5 on both pectoral fins.

Linear measurements can be impacted by allometry. To test for the effect of allometry on our data, the slopes of each trait *versus* standard length were compared between treatments using the package emmeans v1.9.0 [46]. All linear measures followed consistent allometry trends between treatments. Therefore, size-corrected values of each measurement were calculated using standard length (*i.e.*, Trait Ratio = Trait Length/Standard Length). In this study, we prioritize presentation of size-corrected values, but all major results are consistent when both raw measurements and size-corrected values are analyzed.

To test the effect of the experimental treatment on fin ray count and linear measures, we compared the group means of each measurement using an ANOVA (aov), and *post-hoc* pairwise comparisons of groups means were evaluated with Tukey’s Honest Significant Difference tests (Tukey’s HSD). Lastly, to test for the effect of treatment on the variance of the traits, we used the function var.test [47], which uses a standard F-test to calculate differences in group variation. All tests were performed in the R v4.3.2 environment [47].

### Fin ray branching

Analyses of radial morphology indicated plasticity in the anterior portion of the pectoral fin endoskeleton, therefore we tested for differences in branching patterns among treatments in the anterior fin rays. Specifically, we analyzed rays 3-5, which articulate upon radial 1. Rays 1 and 2, which form the leading edge of the fin, are unbranched. Rays 3-5 each can show a single bifurcation. We recorded the presence or absence of branching in each ray and then compared the frequency of branching for each ray between treatments. Lastly, we tested for symmetry in the branching occurrence between left and right pectoral fins. To accomplish this, we created contingency tables that reflected a binary value (*e.g.*, if ray 3 branched on both left and right fins, this value was a 1, if not the value was a 0) and then subjected the frequency of values within the different treatments to a χ^2^ analysis test through the package kim [48]. All tests were performed in the R v4.3.2 environment [47].

To contextualize analyses of branching and inform hypotheses of typical fin ray growth, we analyzed pectoral fin ray branching patterns of wild-caught *S. daemon* from Smithsonian’s Fishes Collection (USNM 3361, n=12; USNM 257536, n = 2). We photographed the left pectoral fins of these individuals, counted the number of branches in pectoral fin ray 3, and recorded standard length of the specimens.

### Geometric morphometrics

Both radial morphometrics and analyses of fin ray branching indicated plasticity in the anterior portion of the pectoral fin. Therefore, to test whether the morphology of fin ray attachment sites can show a plastic response, we analyzed the shape of the base of ray 3 [49]. Two-dimensional cartesian coordinate data were collected using Stereomorph v1.6.7 [50] in R [47]. In total, we used nine landmarks, four fixed and five semi-landmarks to capture the internal, posterior, and dorsal processes of the dorsal hemitrichia. X,Y coordinate data were subjected to generalized Procrustes analysis [51] utilizing bending energy. We then used geomorph v4.0.6 [52–54] to analyze allometry (procD.lm; plot.allometry), and compare across-treatment variation (morphol.disp), and we used the package RRPP v1.4.0 [52] to make pairwise comparisons from the procD.lm output of group means. To aid in data visualization, we conducted a principal component analysis (PCA) and projected the first two components.

### Micro-computed tomography of fin rays

To test for plasticity in the cross-sectional morphology and dorsoventral patterning of pectoral fin rays, dissected pectoral fins were µCT scanned at The University of Chicago, Department of Organismal Biology and Anatomy PaleoCT scanning facility on a GE Phoenix vjtomejx 240 kv/180 kv scanner (PaleoCT facility, RRID:SCR024763, University of Chicago) (pelagic, n=11; rock, n=9; sand, n=8). Scan parameters are as follows: 50kV, 220µA, 150ms, with 1000 projections, and a frame averaging of 3. Voxel sizes are presented in Supplementary Table 1. µCT data were reconstructed with Phoenix Datosjx 2 (v2.3.3) and imported to VGStudio Max (v2.2) to be exported as a tiff stack. µCT data were manually segmented in Avizo v2021.2 (Thermo Fisher Scientific).

To test for differences in cross-sectional area (CSA), we digitally dissected the second, third, and fourth segments of ray 3 and compared CSA between treatments. As with the morphometric analysis, described above, ray 3 was selected for study because radial morphometrics and fin ray branching data suggested plasticity in this part of the fin. After segmentation in Avizo, rays were exported as a tiff stack and converted to binary using the Huang method in ImageJ [45]. We used the function “Moments of Inertia” from the package BoneJ v7.0.18 [55] in ImageJ [45] to calculate CSA of rays from cross-sections oriented orthogonal to the long axis of the ray. The mean CSA of each ray was calculated, size standardized, and compared between treatments. Measurements of CSA were analyzed in R [47] using ANOVA (aov), followed by a Tukey’s Honest Significant Difference test (Tukey’s HSD), and the var.test function in R [47] to test for differences in variation between treatments.

To test for differences in dorsoventral patterning of the fin rays, the dorsal and ventral hemitrichia of ray 3 were segmented in Avizo and compared to one another. For each hemitrichia, CSA and the second moment of area in the major axis (SMoA_maj_), which is a functional measure of the hemitrichia that corresponds to its resistance to bending in the dorsoventral direction, were calculated following methods described above for the full fin rays. For both CSA and SMoA_maj,_ we subjected the dorsal and ventral hemitrichia to both ANOVA and Tukey’s tests with a linear model that allowed us to determine whether hemitrichia are symmetrical and the effect of treatment on symmetry: Variable ∼ Side * Treatment, where Side corresponds to whether a hemitrichia is from the dorsal or ventral side. We also tested whether the magnitude of asymmetry between hemitrichia differed between treatments by calculating the ratio of dorsal/ventral CSA and SMoA_maj_ for each ray and comparing the distribution of these values between treatments using an ANOVA (aov) in R [47].

## RESULTS

### Treatments affect anteroposterior patterning of radials

The length of radial one differs between treatments, and more posterior radials do not show differences in length between treatments (Figure 2; Supplementary Table 2). Specifically, rock-winnowing fish have a longer radial 1 than pelagic and sand-winnowing fish. Thus, treatment impacts the degree of anteroposterior patterning of the pectoral fin radials by differentially affecting radial growth across the series. For all radials, treatments show differences in their magnitude of variation in radial length (Figure 2*b-e*, Supplementary Table 2). Uniformly, the rock-winnowing fish exhibit greater variance than pelagic and sand-winnowing fish. We found no significant differences in radial widths between treatments and no differences in their magnitude of variance of radial widths (Supplementary Table 3).

**Figure 2.**
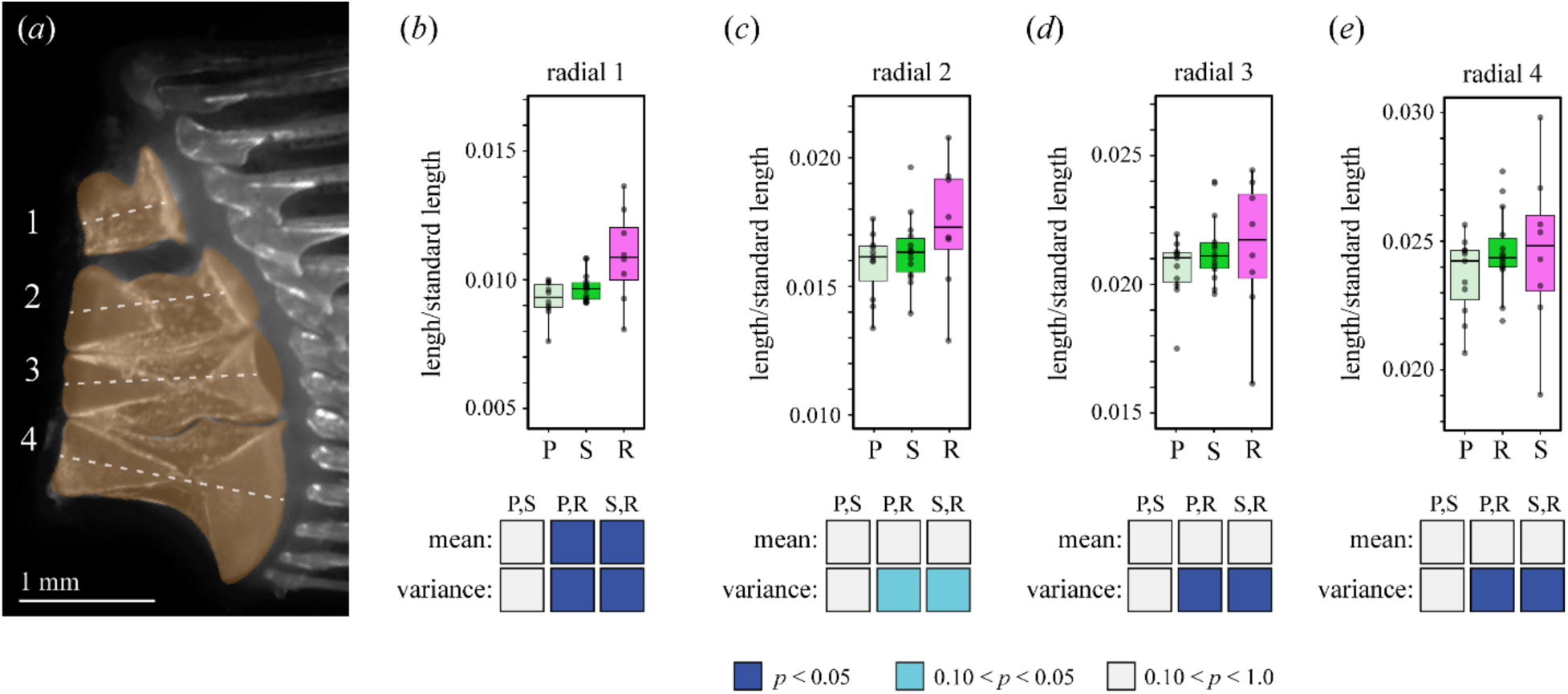
Treatments affect anteroposterior patterning and variation in radial length. (*a*) Photograph of a cleared and stained left pectoral fin with color overlay. Fins were dissected, radial lengths were measured, size corrected by standard length, and compared across treatments. (*b*) In rock-winnowing fish, radial 1 is longer than in other treatments. (*c-e*) Radials 2-4 do not differ in mean length between treatments. For all radials, rock-winnowing fish show greater variation as compared to pelagic and sand-winnowing fish. Matrices reporting P values of group pairwise comparisons of means and variance presented below. Associated statistics reported in Supplementary Table 1. Box plots show median value, the first and third quartiles, and maximum and minimum values. (Abbreviations: P, pelagic treatment; R, rock-winnowing treatment; S, sand-winnowing treatment)

### Treatments affect anteroposterior patterning of fin ray branching

Patterns of fin ray branching differed between treatments (Figure 3*a*). Individuals from the pelagic treatments showed the greatest frequency of branching, and those from the rock-winnowing treatment showed the lowest occurrence of branching (Figure 3*a*,*b*). χ^2^ tests showed significant differences for the left pectoral fin for rays 3 and 4 (P = 0.055 and 0.036, respectively), but not for ray 5 (P = 0.1581). *Post-hoc* tests showed significant differences between rock-winnowing fish and pelagic fish for rays 3 and 4 (P = 0.027 and 0.056, respectively; Supplementary Table 4). In the right pectoral fin, broad differences in the frequency of ray branching between treatments hold: branching is most common in pelagic fish and least common in rock-winnowing fish; however, we found no significant *a priori* χ^2^ comparisons for ray 3,4, or 5 (P = 0.21, 0.24, and 0.51).

**Figure 3.**
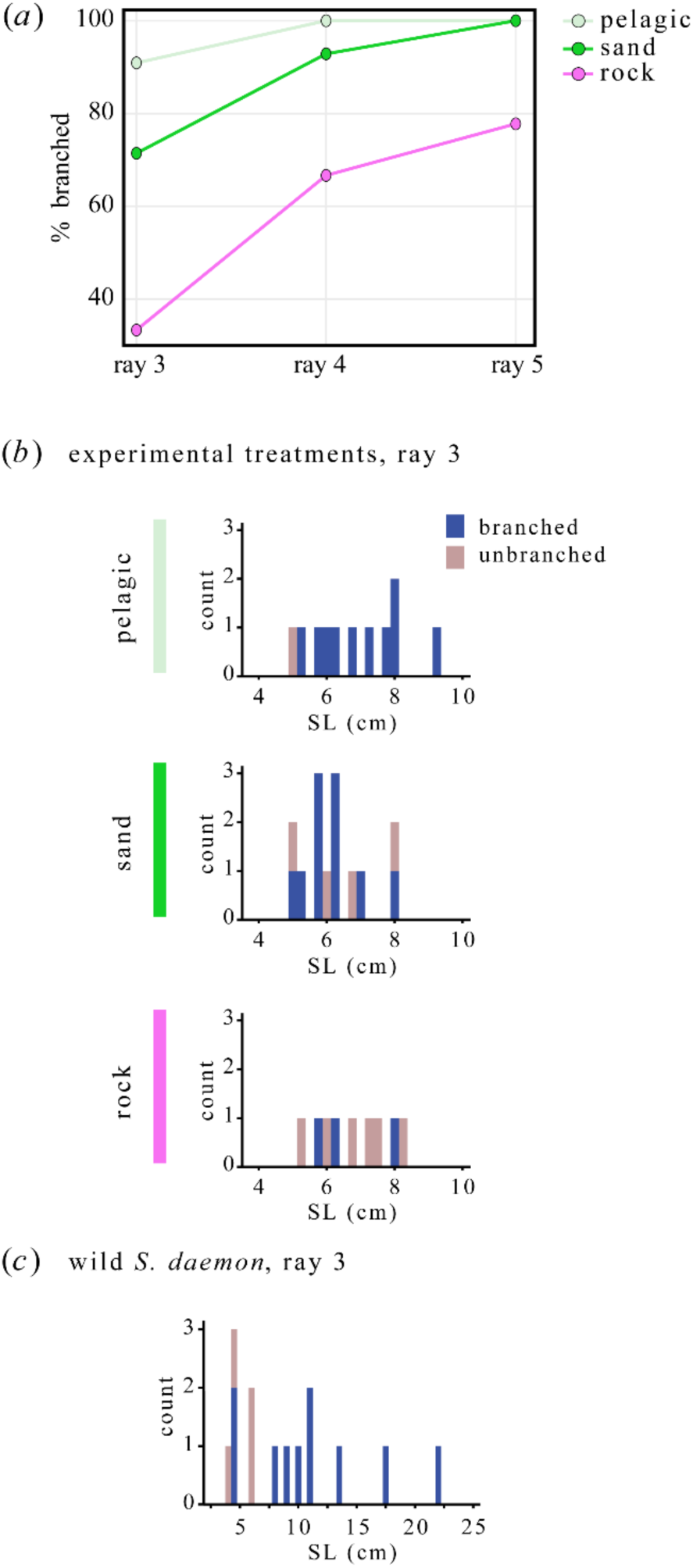
Treatments affect anteroposterior patterning of fin ray branching. (*a*) The frequency of branching in rays 3-5, the three anterior-most branching rays, in experimental populations. For each of the rays, pelagic fish show the greatest frequency and rock-winnowing fish show least frequency of branching. The magnitude of difference between treatments is greatest in ray 3 and decreases more posteriorly. Associated statistics shown in Supplementary Table 3. (*b*) Histogram showing the number of individuals with branched or unbranched ray 3 by standard length for each treatment. (*c*) Same counts as panel *b* for wild-caught *S. daemon* showing that fin ray branching is uniformly present in individuals larger than 8 cm SL. All data are for left pectoral fins.

Because we observed statistical differences between the left and right pectoral fins in their branching patterns, we tested whether treatments differed in tendency to be symmetrical in their fin ray branching. *A priori* χ^2^ tests find that the frequency of asymmetry in branching patterns trends toward significance between treatments for ray 3 (P = 0.067), but not rays 4 and 5 (P = 0.247 and 0.111, respectively). *Post-hoc* tests suggest a difference between rock-winnowing fish and pelagic fish for ray 3, with this comparison approaching significance (P = 0.093). Thus, treatment might affect canalization of fin ray branching between left and right sides of the body, although statistical support is weak.

To assess normal patterns of fin ray growth and branching, we studied wild-caught *S. daemon*. In wild fish, the presence of branching in pectoral fin ray 3 is variable in smaller individuals, and the ray is uniformly branched in individuals larger than 8 cm SL (Figure 3*c*). Therefore, differences in the branching of anterior rays in *S. daemon* under the three experimental treatments might reflect differences in the size at which branching is initiated.

### Fin ray base shape is robust to experimental treatments

To test for differences in the shape fin ray muscle attachment sites, we used four fixed landmarks and five semi-landmarks across the base of ray three (Supplemental Figure 1*a*). Tests of allometry revealed similar parallel allometries between the three treatments and no significant effect of size on these data (P = 0.7158, R^2^ = 0.013) (Supplemental Figure 1*b*). Allometry is, therefore, unlikely to affect comparisons of shape between treatments, and size was therefore not treated as a covariable in subsequent analyses. A PCA revealed that the first two axes of variation contained ∼81% of the variation in our data. Convex hulls indicate that pelagic fish occupied the greatest area of morphospace and that rock-winnowing fish are restricted along PC1 as compared to other treatments (Supplemental Figure 2*c*). Despite deformation grid projections suggesting disparate shapes between rock-winnowing fish and the other two groups (Supplemental Figure 1*d*), we found no significant differences between group means (Supplemental Figure 1*e*). Likewise, tests of morphological disparity revealed no significant differences among group variation; however, comparisons of rock-winnowing and sand-winnowing fish recovered P values that approached significance when compared to the pelagic fish (P = 0.1055 and P = 0.0757, respectively) (Supplemental Figure 1*e*).

### Rock-winnowing fish have greater variation in cross-sectional area of fin rays

Differences in fin ray branching were most pronounced in ray 3 of the pectoral fin; therefore, we tested for differences in cross-sectional morphology of this ray (Figure 4*a*). Treatments did not affect mean cross-sectional area (CSA) (P = 0.513). However, statistics from F-tests reveal trending differences in variance between groups with the rock-winnowing fish having greater variance than pelagic (P = 0.074) and sand-winnowing fish (P = 0.031) (Figure 4*b*). There was no difference in variance between pelagic and sand-winnowing fish (P = 0.737).

**Figure 4.**
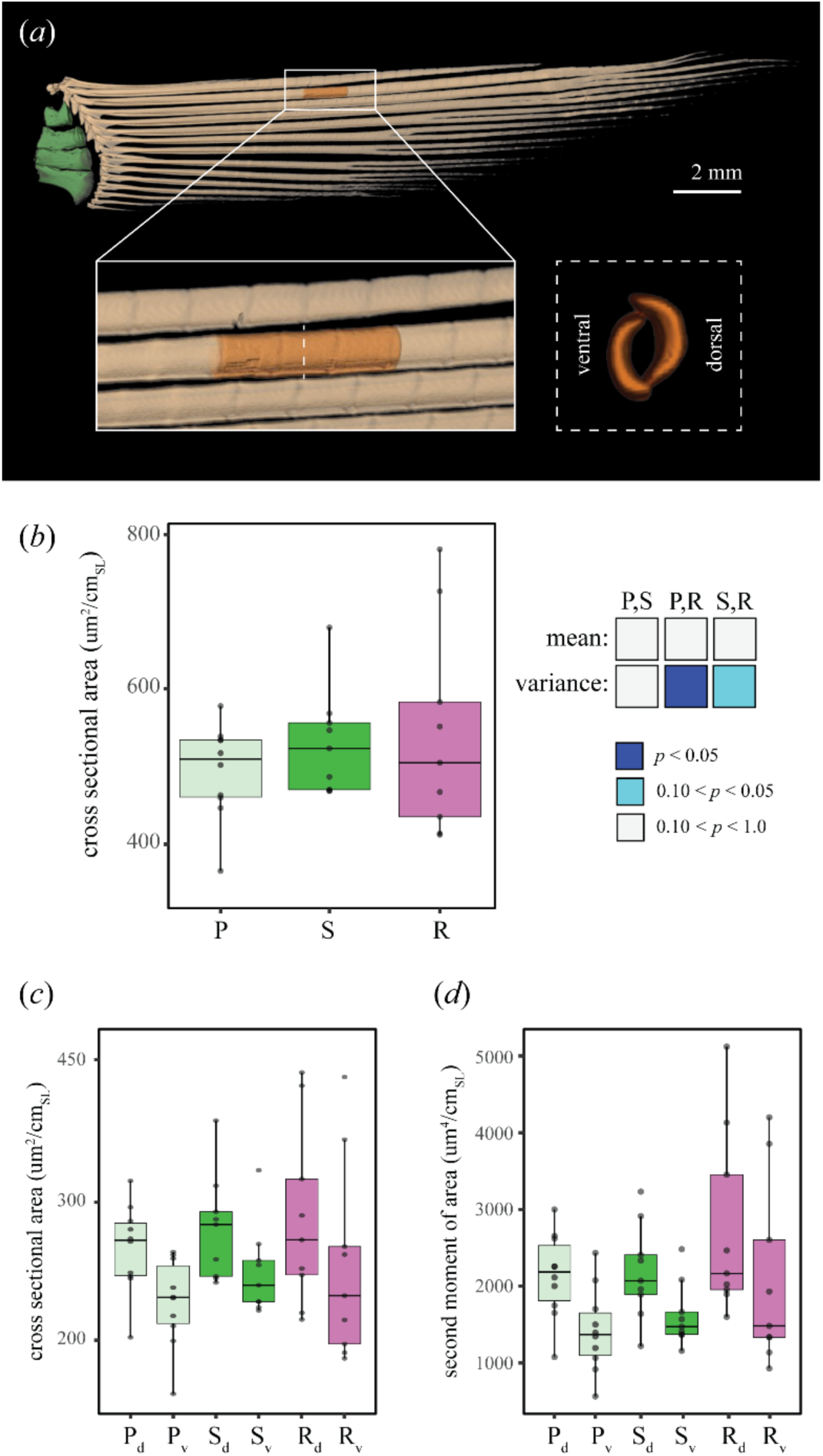
Treatments differ in variance of CSA but not in dorsoventral patterning. (*a*) Volumetric rendering of μCT scan of a pectoral fin with insert showing the three segments that were analyzed. (*b*) The mean CSA of the segments 2-4 on ray 3 does not differ between treatments, but rock-winnowing fish have greater variance in CSA than either sand-winnowing or pelagic treatments. (*c*) CSA of dorsal and ventral hemitrichia across the three treatment groups. (*d*) Second moment of area along the major axis for both dorsal and ventral hemitrichia across the three treatment groups. Box plots show median value, the first and third quartiles, and maximum and minimum values. (Abbreviations: P, pelagic treatment; R, rock-winnowing treatment; S, sand-winnowing treatment)

We tested whether treatment affected dorsoventral patterning by comparing CSA and SMoA_maj_ of the dorsal and ventral hemitrichia of ray 3 using the model: Trait ∼ Side * Treatment. We find statistically significant asymmetries in dorsal and ventral hemitrichia for both CSA (P = 0.008) and SMoA_maj_ (P = 0.004). In all treatments, dorsal hemitrichia have greater CSA and SMoA_maj_ than ventral hemitrichia (Figure 4*c*, *d*). Next, to test whether treatment affected magnitude of asymmetry of CSA and SMoA_maj_, we calculated an index of asymmetry (dorsal/ventral) of each variable and compared these ratios between treatments with ANOVAs. We find no difference in magnitude of asymmetry between treatments (P = 0.286). Thus, our data do not support the hypothesis that the treatment affects the dorsoventral patterning of fin rays.

### Fin ray segment length is robust to experimental treatments

To test whether the anteroposterior patterning of fin ray segment lengths can reflect plasticity, we measured the first proximal segment in rays 3-5 (Supplemental Figure 2*a*). In the left fin, none of the three rays showed differences in mean lengths between treatments (Figure 6*c-de*), and only one of the pairwise comparisons for differences in magnitude of variation approached significance (Supplemental Figure 2*e*, Supplementary Table 6). In the right pectoral fin, no pairwise comparisons of mean or variance showed differences in length between treatments (Supplementary Table 7). Thus, our data do not support the hypothesis that the treatment affects the anteroposterior patterning of ray segment length.

## DISCUSSION

Our analyses show that altered foraging regime can induce a plastic response in fin skeletal patterning, revealing features of the fin that are either responsive or robust to the environmental pressures assayed here. The anteroposterior patterning of both radials and fin rays are affected, while dorsoventral patterning appears unaffected. Some features, like the cross-sectional area of fin rays, indicate altered patterns of variability, while others, like the lengths of fin ray segments, do not show differences between the treatments. Collectively, these data suggest that plasticity contributes to the morphological disparity of fins and implicates hydrodynamic loading during ontogeny as a factor that affects the development and canalization of fin skeletons.

### Behavior and swimming alter fin architecture

Observed phenotypic differences between treatments likely reflect altered patterns of pectoral fins use when fish were induced to forage in different ways. Although this study did not interrogate kinematics, observations during the experimental treatment indicate that in *S. daemon* the angle of approach differs when feeding in open water as compared to when winnowing (*i.e.*, head directed upwards *vs* downwards), and that sediment granule size also likely affects body positioning during sediment uptake and expulsion during winnowing (Supplementary Movies 1-3). Thus, we predict that different swimming patterns result in distinct hydrodynamic loads experienced by the fins and that this affects patterns of bone growth. We regard this as more likely than alternative hypotheses, for example of pre-existing differences between groups, as individuals were simultaneously collected from the same locality, or of differences in growth rate or allometry between treatments, as individuals were of consistent sizes following the five months of treatment (Figure 3*b*), and because our results are consistent when both raw measurements and size-corrected values are analyzed.

Altered swimming behaviors are likely responsible for differences in the anteroposterior patterning of both the endoskeleton and dermal skeleton. In the pectoral fin, mean differences in trait values were observed in anterior elements (*e.g.*, length of radial 1 and branching of fin ray 3), while more posterior elements were less likely to show consistent differences between treatments (*e.g.*, lengths of radials 2-4 and branching of fin rays 4 and 5). This anteroposterior polarity is consistent with the hypothesis that differences are mechanically induced. Hydrodynamic loads are not uniformly applied to fins during swimming, as the leading edge of flapping foils experience higher magnitudes of loading as compared to the trailing edge [18], and in skeletal systems those areas of highest magnitude loading are more prone to plastic response [56].

In response to our experimental treatments, some traits showed differences in their mean values, while others differed in the magnitude of variance around the mean trait value. We predict that the traits that showing changes in mean value (*e.g.*, rock-winnowing fish having a longer radial 1 and being more likely to have an unbranched ray 3 of the pectoral fin) reflect differences in mean behavior by treatment. By contrast, those traits that show different patterns of variability (*e.g.*, rock-winnowing fish having greater variability in the lengths of all radials and in fin ray CSA as compared to other treatments, or rock-winnowing fish having greater asymmetry in fin ray branching between left and right fins) are likely related to *S. daemon* exhibiting greater kinematic variability under some regimes. Regardless of the underlying behavioral dynamics, the observed differences in phenotypic variability reveal how environmental pressures, like how food resources are distributed within an environment, can impact trait canalization and reveal fodder upon which natural selection can act [57–59].

### Potential developmental mechanisms for plasticity in the fin skeleton

The anteroposterior patterning of fins has previously been interrogated through genetic approaches. These developmental genetic studies have yielded numerous factors that contribute to adult fin morphology and to the evolution of fin diversity [60–63]. To our knowledge, our results are the first to implicate swimming and hydrodynamic loading on anteroposterior patterning of fins, raising questions of how early patterning processes and plastic responses over ontogeny interact to produce disparity in fish fins. For example, in our study, observed differences in radial length manifest in juvenile fishes, after early developmental patterning occurs [34,60–62,64,65] and likely represent a separate mechanism of regulating radial length.

Treatments differed in the likelihood of bifurcations of the anterior rays in the pectoral fin. The boundary between unbranched and branching rays is a typical measure of fin patterning [41,66]. Numerous factors are known to determine patterns of fin ray branching, including local *Shh* expression [32,67], thyroid hormone signaling [68], inter-ray blastemas [69], and osteolytic tubules [70]. Given that the *Shh* and thyroid pathways are sensitive to mechanical and environmental inputs [71–75], these are strong candidates for the mechanism behind differences in branching between treatments in our study. More generally, however, our results suggest that loading of the distal fin during fin ray development contributes to the initiation of fin ray bifurcation. How the fin rays of *S. daemon* detect the loading warrants further study. Perhaps it is by a *sost/*sclerostin-dependent mechanism [76,77], which is indeed expressed in anosteocytic bone and has been linked to mechanical loading and bone remodeling and repair. Or, this could be through another mechanism, such as by the primary cilia in the surrounding tissues, which are known mechanoreceptors that play a key role in bone remodeling [78–80]. Those two possibilities are not mutually exclusive and could be connected.

Several fin traits did not change in response to our treatments. These features are either fixed before treatments began, unable to vary response to differential loading regimes, or our experiments did not reach thresholds to induce responses. A previous study in African cichlids found that altered feeding regime affected the number of pectoral fin rays [38]; however, we did not observe altered ray counts in our analysis. Fin ray number has been linked to variation in the expression of wnt7aa, lef1, and col1a1 signaling, which occurs as early as 6 days post-fertilization in developing African cichlids [33], a timepoint significantly prior to the beginning of our experiment. Perhaps our treatments began after fin ray number is fixed in *S. daemon*. In *S. daemon*, the length of the first fin ray segment did not show plastic response. This is consistent with past work that predicts fin rays grow by distal addition to the rays rather than changes in segment length [81] and studies of caudal fin regeneration in zebrafish [82], which suggest the fin ray segmentation system operates by mechanisms independent of the of fin loading during outgrowth.

Finally, our experiments did not show altered patterns of dorsoventral patterning in the fin rays. As compared to other fin patterning axes, dorsoventral patterning has received significantly less attention [41,83]. Perhaps differences in loading in our treatment did not reach thresholds to produce altered hemitrichia size or shape, and it is worth interrogating whether altered dorsoventral patterning changes can be induced in other contexts, like walking fishes on terrestrial environments, where fin rays must contend the effect of gravity [37,41,84,85].

### The functional and evolutionary implications of fin plasticity

Across fish species, the form and function of fins are tightly linked [18,20–23,27,28]. We predict that phenotypic differences between our treatments translate to and result from differences in fin function. For example, the length of radial 1 is greater in rock-winnowing fishes as compared to pelagic fishes. The muscles involved in raising, lowering, and abducting the pectoral fin originate from the pectoral girdle, span the radials, and attach to the base of the fin rays (Winterbottom 1973). Differences in radial length should, therefore, correspond to differences in mechanical advantage of the system (*e.g.*, differences in the relative speed and force production during up and downstrokes of the fin).

Additionally, differences in fin ray branching should impact the effective flexural stiffness of the fin web [83]. Rock-winnowing fishes are less likely to have branched anterior fin rays as compared to pelagic fishes, which implies a lower effective flexural stiffness in the leading edge of the fin [83,86]. A comparatively reduced effective flexural stiffness could contribute to enhanced maneuverability as, in pelagic fishes, increasing flexural stiffness is associated with enhanced mechanical efficiency and decreased stiffness is associated with enhanced maneuverability [18,23,87].

These data implicate juvenile foraging behaviors as a potential cause of inter-specific differences in fin morphology and the adaptive diversification of fishes. Previous work on fin plasticity has primarily considered major differences in environment (*e.g.*, aquatic *vs* semi-terrestrial conditions; natural riverine systems *vs* captive tank-raised fishes) and implicated plasticity in major ecological transitions [37] and the origin of novel skeletal traits [39]. Our treatments differed in more subtle ways, varying whether food was presented in the water column or in sandy or rocky substrate. Differences in substrate type are common across aquatic environments, and can result from anthropogenic causes, like dam construction [88]. The phenotypic responses observed in our study might, therefore, be within the range of environmental pressures that fishes are likely to confront in the wild and contribute to fin disparity in wild fishes.

Several of our results suggest our sand-winnowing treatment to be intermediate when compared to both rock-winnowing and pelagic feeding treatments. The intermediate nature of sand-winnowing matches our previous study in craniofacial anatomy of these same specimens [44], which suggested that several craniofacial traits of the sand-winnowing population were the “average”, with rock and pelagic foraging fish being on the opposite extremes of our sampling. This is especially notable when considering that the sandy substrate environment is most typical for what *S. daemon* would encounter [is there a citation for this? Ok if not], leaving a forced pelagic or rocky environment to be considered novel foraging environments.

Our study provides evidence that foraging behaviors in fishes can alter both craniofacial morphology [44] and fin patterning. This is consistent with previous work in cichlids [38,44]. In recent years, strides have been made in testing hypotheses of evolutionary integration between craniofacial and fin systems, supporting long standing eco-morphological predictions that they are linked [27,28,89]. However, the role that plasticity can play in placing crania and fins on the integration-modularity spectrum is still unknown, and future work should test how plasticity impacts correlations between the craniofacial and locomotor systems. Such experimental studies can reveal which systems of the Baupläne are integrated or modular and clarify the forces that impact the development and evolution of fins in fishes, broadly.

## ACKOWLEDGEMENTS

We thank Diane Pitassy and Kris Murphy at the Smithsonian for their support and for providing us with access to the collections. We thank Skylar Mannen for her assistance in data recording and editorial feedback during the early writing process. Funding by the Natural History Collections at the University of Massachusetts, and The Pennsylvania State University Department of Biology and Erickson Discovery Award. Yara Haridy was supported by funds awarded to N.H. Shubin from the Brinson Family Foundation and the Biological Sciences Division of The University of Chicago.

## DATA AVAILAILITY

All µCT data are available for download from Morphosource (DOI provided upon acceptance). Linear measure data will be made available on GitHub at https://github.com/DrMermaid-MichelleGilbert (stable link provided upon acceptance).

---

**Supplemental Figure 1.**
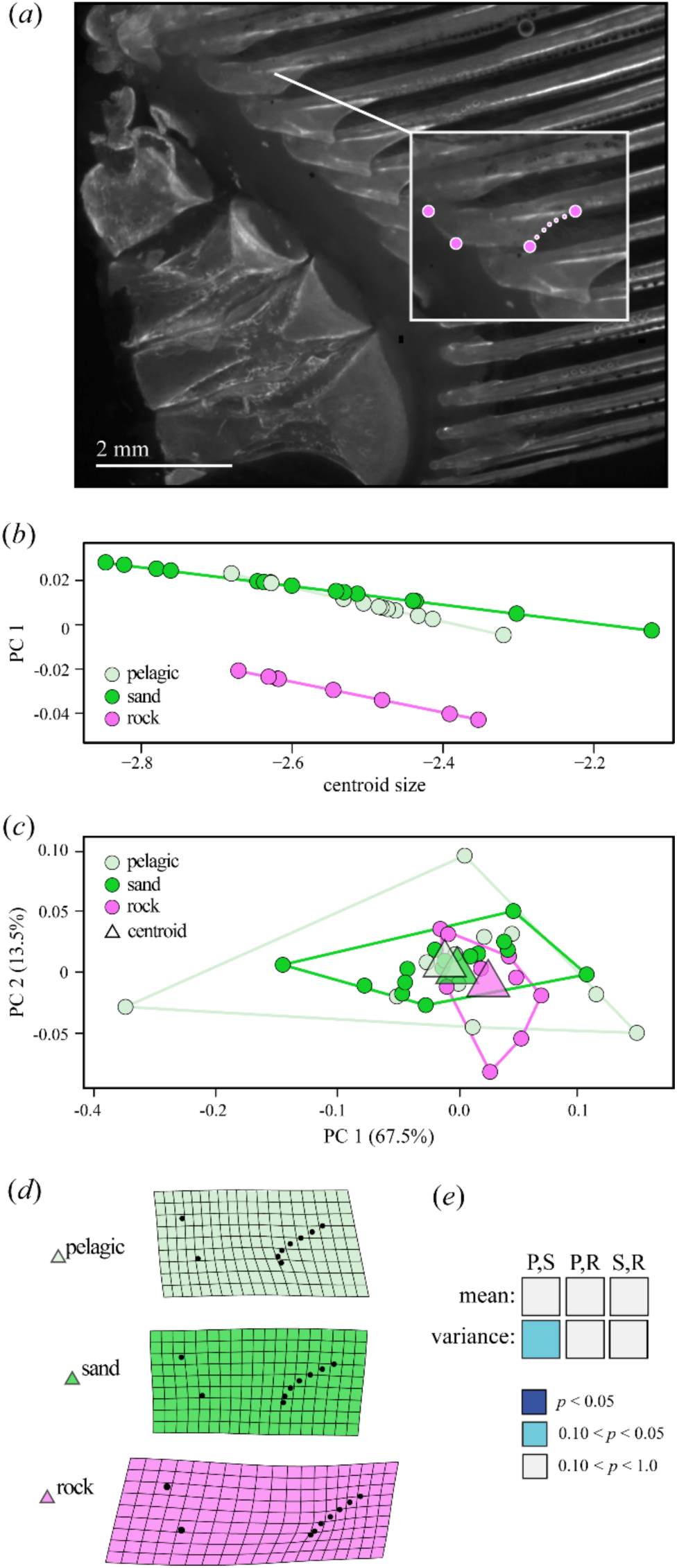
Treatments do not clearly affect shape of the fin ray base. (*a*) Photograph of a dissected cleared and stained left pectoral fin. Insert represents landmark configuration used to quantify the shape of the muscle attachment site on ray 3: large circles are fixed landmarks; small circles are semi-landmarks. (*b*) Regression of the first primary axis of variation against centroid size, demonstrating parallel allometric trajectories. (*c*) Principal component plot of the first two axis of variation. Circles represent individuals, triangles represent group means. (*d*) Mean deformation grids illustrating the three treatment groups. Differences magnified by a factor of 3. (*e*) Matrix reporting P values of group pairwise comparisons of both means and variance. (Abbreviations: P, pelagic treatment; R, rock-winnowing treatment; S, sand-winnowing treatment)

**Supplemental Figure 2:**
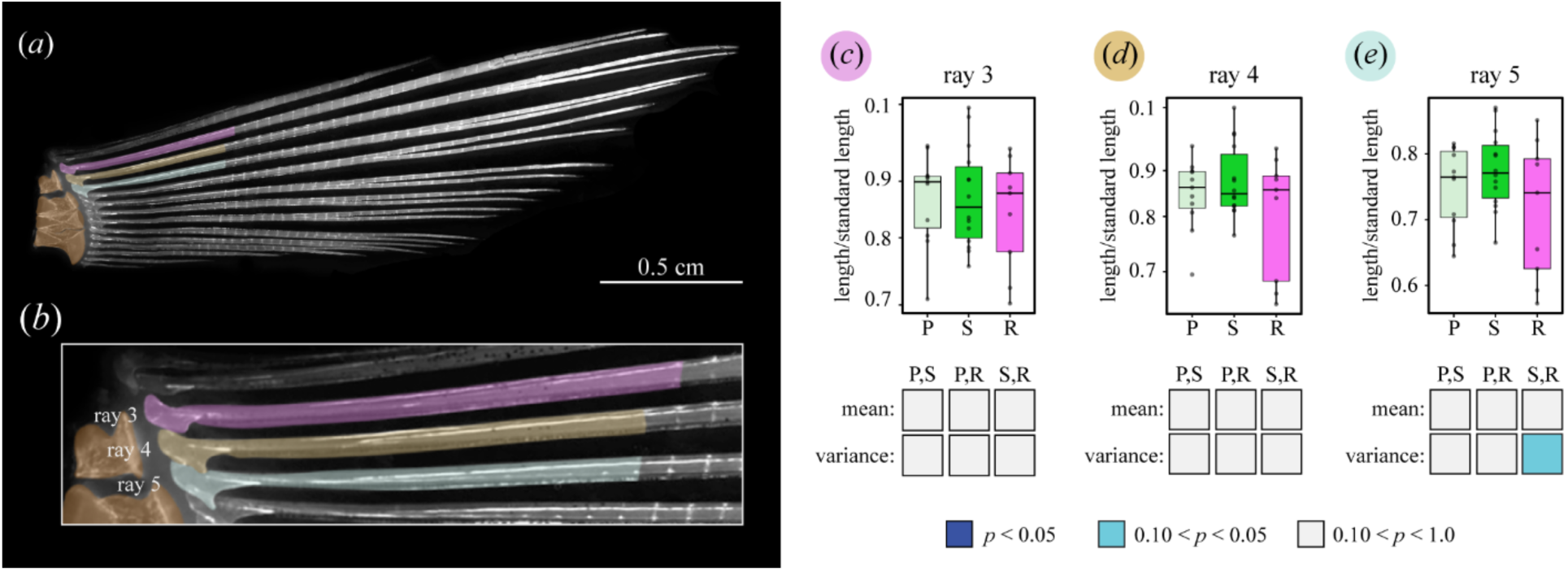
Treatments do not affect fin ray segment length. (a, *b*) Photographs of a cleared and stained pectoral fin with color overlay showing the rays and segments that were measured. (*c-d*) The mean length of the proximal fin ray segment does not differ between treatments. For one of the rays, ray 5, one of the comparisons approaches significance threshold of P < 0.05. Associated statistics shown in Supplementary Table 6. Box plots show median value, the first and third quartiles, and maximum and minimum values. (Abbreviations: P, pelagic treatment; R, rock-winnowing treatment; S, sand-winnowing treatment)

**Supplementary Table 1:** Voxel sizes from µCT scans of *Satanoperca daemon.* All other scan parameters are presented in the main text

| specimen ID | treatment | voxel size (μm) |
| --- | --- | --- |
| 1 *S.daemon* *Fin* *PelagicTr* | Pelagic | 11.345 |
| 2 *S.daemon* *Fin* *PelagicTr* | Pelagic | 11.942 |
| 4 *S.daemon* *PelagicTr* | Pelagic | 13.019 |
| 5 *S.daemon* *Fin* *PelagicTr* | Pelagic | 15.144 |
| 6 *S.daemon* *Fin* *PelagicTr* | Pelagic | 10.71 |
| 8 *S.daemon* *Fin* *PelagicTr* | Pelagic | 13.861 |
| 9 *S.daemon* *Fin* *PelagicTr* | Pelagic | 13.757 |
| 10 *S.daemon* *Fin* *PelagicTr* | Pelagic | 9.895 |
| 11 *S.daemon* *Fin* *PelagicTr* | Pelagic | 14.964 |
| 13 *S.daemon* *Fin* *PelagicTr* | Pelagic | 11.353 |
| 13 *S.daemon* *Fin* *PelagicTr* | Pelagic | 11.248 |
| 1 *S.daemon* *Fin* *RockTr* | Rock | 11.304 |
| 1 *S.jurupari* *Fin* *RockTr* | Rock | 11.353 |
| 2 *S.daemon* *Fin* *RockTr* | Rock | 12.721 |
| 3 *S.daemon* *Fin* *RockTr* | Rock | 10.786 |
| 4 *S.daemon* *Fin* *RockTr* | Rock | 9.288 |
| 6 *S.daemon* *Fin* *RockTr* | Rock | 11.353 |
| 7 *S.daemon* *Fin* *RockTr* | Rock | 11.353 |
| 9 *S.daemon* *Fin* *RockTr* | Rock | 11.353 |
| 12 *S.daemon* *Fin* *RockTr* | Rock | 12.361 |
| 3 *S.Daemon* *Fin* *SandTr* | Sand | 14.699 |
| 4 *S.Daemon* *Fin* *SandTr* | Sand | 14.6 |
| 9 *S.daemon* *Fin* *SandTr* | Sand | 12.361 |
| 11 *S.daemon* *Fin* *SandTr* | Sand | 11.134 |
| 15 *S.daemon* *Fin* *SandTr* | Sand | 11.134 |
| 16 *S.daemon* *Fin* *SandTr* | Sand | 10.213 |
| 17 *S.daemon* *Fin* *SandTr* | Sand | 10.213 |
| 18 *S.daemon* *Fin* *SandTr* | Sand | 10.213 |

**Supplementary Table 2:** Statistics for comparisons of radial length. P values are reported from Tukey’s HSD test. Comparisons of mean are below the diagonal, and comparisons of variance are above the diagonal. P values < 0.05 are shown in bold. P values approaching the threshold of 0.05 are italicized. (Abbreviations: P, pelagic treatment; R, rock-winnowing treatment; S, sand-winnowing treatment)

|  | radial 1 |  |  | radial 2 |  |  | radial 3 |  |  | radial 4 |  |  |
| --- | --- | --- | --- | --- | --- | --- | --- | --- | --- | --- | --- | --- |
|  | P | R | S | P | R | S | P | R | S | P | R | S |
| P | - | **0.014** | 0.4396 | - | *0.0586* | 0.9076 | - | **0.0217** | 0.7658 | - | **0.03568** | 0.9533 |
| R | **0.0056** | - | **0.0005** | 0.143 | - | *0.0521* | 0.5573 | - | **0.0257** | 0.5612 | - | **0.0257** |
| S | 0.5348 | **0.0325** | - | 0.7129 | 0.3925 | - | 0.5033 | 0.9977 | - | 0.4657 | 0.9999 | - |

**Supplementary Table 3:**
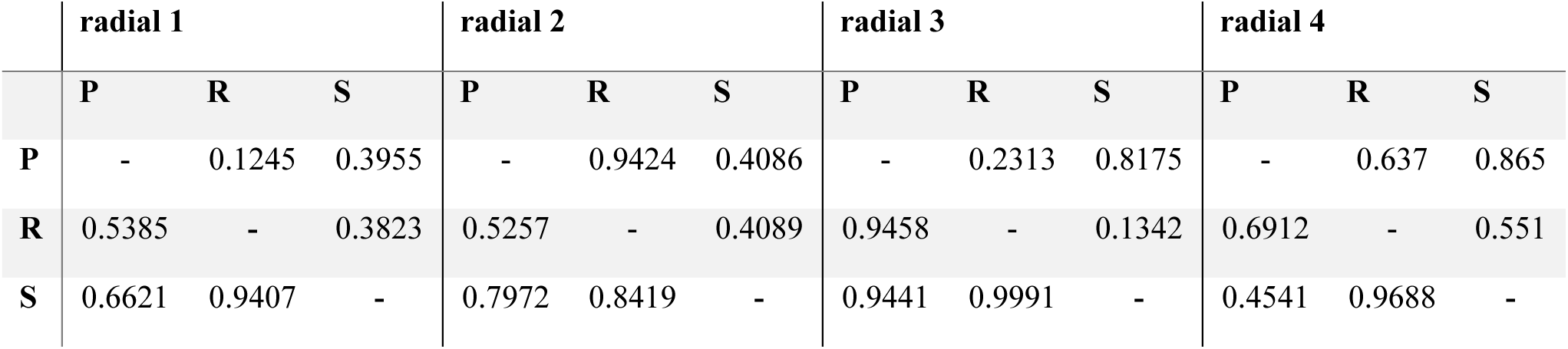
Statistics for comparisons of radial width. Differences in radial width are not detected between the three treatments. P values are reported from Tukey’s HSD test. Comparisons of mean are below the diagonal, and comparisons of variance are above the diagonal. (Abbreviations: P, pelagic treatment; R, rock-winnowing treatment; S, sand-winnowing treatment)

**Supplementary Table 4:** χ^2^ statistics for comparisons of left pectoral fin branch numbers. P values are reported for rays 3 and 4, where an *a priori* test shows significance. P values < 0.05 are shown in bold. P values approaching the threshold of 0.05 are italicized. (Abbreviations: P, pelagic treatment; R, rock-winnowing treatment; S, sand-winnowing treatment)

|  | *ray 3* |  | *ray 4* |  | *ray 5* |  |
| --- | --- | --- | --- | --- | --- | --- |
|  | **P** | **R** | **P** | **R** | **P** | **R** |
| **R** | **0.027** | - | *0.056* | - |  | - |
| **S** | 0.481 | 0.171 | 1 | 0.110 |  |  |

**Supplementary Table 6:** Statistics for comparisons of proximal segment length from the left pectoral fin. Statistics reported from Tukey’s HSD test. Comparisons of mean are below the diagonal, and comparisons of variance are above the diagonal. P values approaching the threshold of 0.05 are italicized. (Abbreviations: P, pelagic treatment; R, rock-winnowing treatment; S, sand-winnowing treatment)

|  | proximal segment |  |  |
| --- | --- | --- | --- |
| ray 3 | P | S | R |
| P | - | 0.7786 | 0.535 |
| S | 0.9926 | - | 0.6941 |
| R | 0.9392 | 0.9290 | - |
| ray 4 | P | S | R |
| P | - | 0.9227 | 0.1113 |
| S | 0.9723 | - | 0.1025 |
| R | 0.7586 | 0.7194 | - |
| ray 5 | P | S | R |
| P | - | 0.8860 | 0.118 |
| S | 0.6958 | - | *0.0681* |
| R | 0.6159 | 0.3646 | - |

**Supplementary Table 7:**
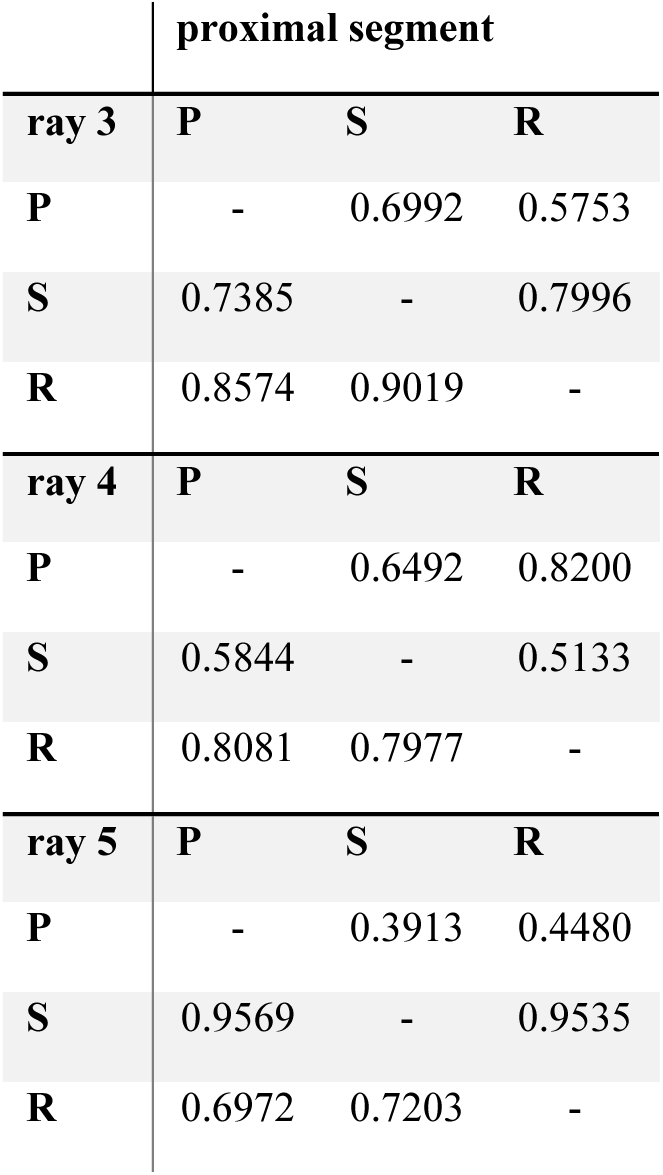
Statistics for comparisons of segment length from the right pectoral fin. Statistics reported from Tukey’s HSD test. Comparisons of mean are below the diagonal, and comparisons of variance are above the diagonal. (Abbreviations: P, pelagic treatment; R, rock-winnowing treatment; S, sand-winnowing treatment)

## Legends for Supplementary Movies

**Supplementary Movie 1:** Representative *S. daemon* feeding bout from the pelagic treatment

**Supplementary Movie 1:** Representative *S. daemon* feeding bout from rock-winnowing treatment

**Supplementary Movie 3:** Representative *S. daemon* feeding bout from the sand-winnowing treatment

**Supplementary Movie 4:** Volumetric rendering of a µCT scan of a pectoral fin of *S. daemon*

